# Nucleosome dynamics and maintenance of epigenetic states of CpG islands

**DOI:** 10.1101/047332

**Authors:** Ian B. Dodd, Kim Sneppen

## Abstract

Methylation in mammalian DNA occurs primarily at CpG sequences. The CpG sites are distributed in high density clusters (or islands) separated by extended regions of low density. Cluster methylation tends to be bimodal, being dominantly unmethylated or mostly methylated. For CpG clusters near promoters, low methylation is associated with transcriptional activity, while high methylation is associated with gene silencing. Alternative CpG methylation states are thought to be stable and heritable, conferring localized epigenetic memory that allows transient signals to create long-lived gene expression states. Positive feedback where methylated CpG sites recruit enzymes that methylate nearby CpGs, does not easily explain that as clusters increase in size or density they change from being primarily methylated to primarily unmethylated. Here, we show that an interaction between the methylation state of a cluster and its occupancy by nucleosomes provides a mechanism to reproduce epigenetic potential and the genome wide systematics of CpG islands.

## I. INTRODUCTION

Methylation of cytosines in the context of CpG sequences in vertebrate DNA is currently of intense interest due to its association with epigenetic gene regulation [1–4]. For alternative CpG methylation states to encode epigenetic memory, these states must be stable and heritable through DNA replication. The symmetry of the CpG sequence on double-stranded DNA provides a simple model for the inheritance of the methylation state of a single CpG [5, 6]. Insertion of unmethylated cytosines during DNA replication means that an unmethylated CpG produces two unmethylated daughter CpGs. A fully methylated CpG site can be inherited if the two hemimethylated daughter CpGs (one C methylated, one unmethylated) produced after its DNA replication are efficiently recognized by maintenance DNA methyltransferases (DNMTs) and converted to fully methylated sites.

However, this ‘classical’ model alone cannot explain a number of more recent observations [7–9]. The high fidelity required for error-free maintenance of methylated sites without unwanted methylation of unmethylated sites is not matched by DNMT enzymes in vitro [9] or in vivo [10]. In addition, further errors can arise from enzymes that are now known to actively demethylate CpG sites [11–14]. Indeed, the frequencies of hemimethylated CpG sites observed by hairpin bisulfite PCR [15] indicate high error rates for individual CpG sites. Variants of the classical model that allow for methylation errors show that, under a given set of conditions, there is only a single equilibrium fractional methylation for any individual CpG, instead of the stable alternative states needed for epigenetic memory [8, 9, 16, 17]. Furthermore, CpG sites display ‘group behaviour’ that is not predicted from models where CpG sites are independent. Clusters of CpG sites in vivo show bimodal methylation, with clusters tending to have either most CpGs methylated (hyper-methylated) or most CpGs unmethylated (hypo-methylated) and avoiding intermediate mixed methylation states [8, 18–22].

As an alternative, we have proposed a model where CpG sites interact with each other, with methylated and hemimethylated CpG sites recruiting DNMTs, and un-methylated CpGs recruiting demethylases, with the recruited enzymes acting to modify other local CpG sites [8]. Simulations show that such a positive feedback can produce an inherently bistable system, allowing CpG sites to collaborate to dynamically maintain either an overall hyper-or hypo-methylated state of a cluster. Thus, bimodal cluster methylation arises naturally from dynamic bistability. Importantly, the hyper-or hypo-methylated states of a CpG cluster could each be inherited over many simulated cell generations, even in the presence of high error rates. Thus, unlike other models, the collaborative model provides for robust epigenetic memory. Bistability of cluster methylation can be seen in vivo, as some CpG clusters are seen in different methylation states in different cells of the same type, and two copies of the same cluster can even be in different methylation states within the same cell [10, 21].

The collaborative model opens for the possibility that the topography of a cluster could influence its methylation state. Indeed, our analysis of the methylation states of CpG clusters in the human genome revealed a strong trend where the probability of hypo-methylation increases with increasing number and density of CpGs in the cluster [22]. Clusters of 10 or fewer CpG sites tend to be hyper-methylated and clusters of 50 or more CpG sites tend to be hypo-methylated, with only intermediate-sized clusters showing bimodal methylation. This behaviour could be reproduced by a model in which methylated CpGs collaborate over longer DNA separations, while unmethylated CpGs collaborate over shorter distances [22]. However, without a satisfying molecular mechanism, these differing distance dependencies in the model are somewhat arbitrary.

A number of observations suggest an interplay between CpG methylation and the nucleosomes that package eukaryotic DNA [7, 23]. The core nucleosome is a complex of 8 histone proteins with ~150 bp of DNA wrapped almost 2 times around it. Nucleosomes are spaced ~170-200 base pairs (bp) apart (centre to centre) across eukaryotic genomes, occupying 75-90% of the DNA [24, 25]. Nucleosome occupancy varies across the genome and is correlated with the local level of DNA methylation [26–28]. Suggested mechanisms for this correlation include a preference by DNMT enzymes to methylate nucleosomal CpGs [26, 28] and exclusion of nucleosomes from unmethylated DNA due to competition with bound transcription factors and other proteins [25].

We show here that interactions between nucleosome occupancy and DNA methylation, whereby unmethylated CpGs inhibit nucleosome occupation and nucleosome occupation stimulates DNA methylation, provide a simple mechanism to generate stable alternative methylation states that are responsive to the size and density of a CpG cluster. Extensions of this basic model that introduce additional collaboration between CpGs increase the stability of the alternative states and improve the match to the behaviour of CpG sites in vivo. The model has few parameters yet can explain overall trends among a majority of the CpG clusters in the human genome as well as provide robust epigenetic memory.

## II. MODEL AND METHOD

This paper scrutinizes the interplay between nucleosome positioning and the methylation status of CpG sites. The model consists of nucleosomes that cover a space of about 140 bp of DNA and make random walks along the DNA, subject to hard core exclusion as in the Tonks gas model [29–32]. The model simulates a finite segment of DNA with open boundaries from which nucleosomes may leave or be inserted. To maintain nucleosome density, nucleosomes are inserted when there is large enough space between adjacent nucleosomes or at the boundaries. In cells, nucleosome movement, insertion and eviction are active processes catalysed by chromatin remodeling and histone chaperone complexes [33]. To keep the model simple, we do not model nucleosome eviction.

Fig. 1 shows a CpG cluster and a portion of the surrounding low CpG density DNA occupied by nucleosomes. The system consists of a constant number *n*_*cpg*_ of sites that are distributed along an *L* bp segment of chromosomal DNA *(L* is much larger than the CpG cluster to avoid external boundary effects on nucleosomes). The DNA is represented in 2 bp units, with each unit a CpG or not. A CpG can be methylated (m), hemimethylated (h) or unmethylated (u). At any time there are *N(t)* nucleosomes on this DNA, and the model is simulated in time units given by 2 bp steps of the random walk of each nucleosome.

**FIG. 1.**
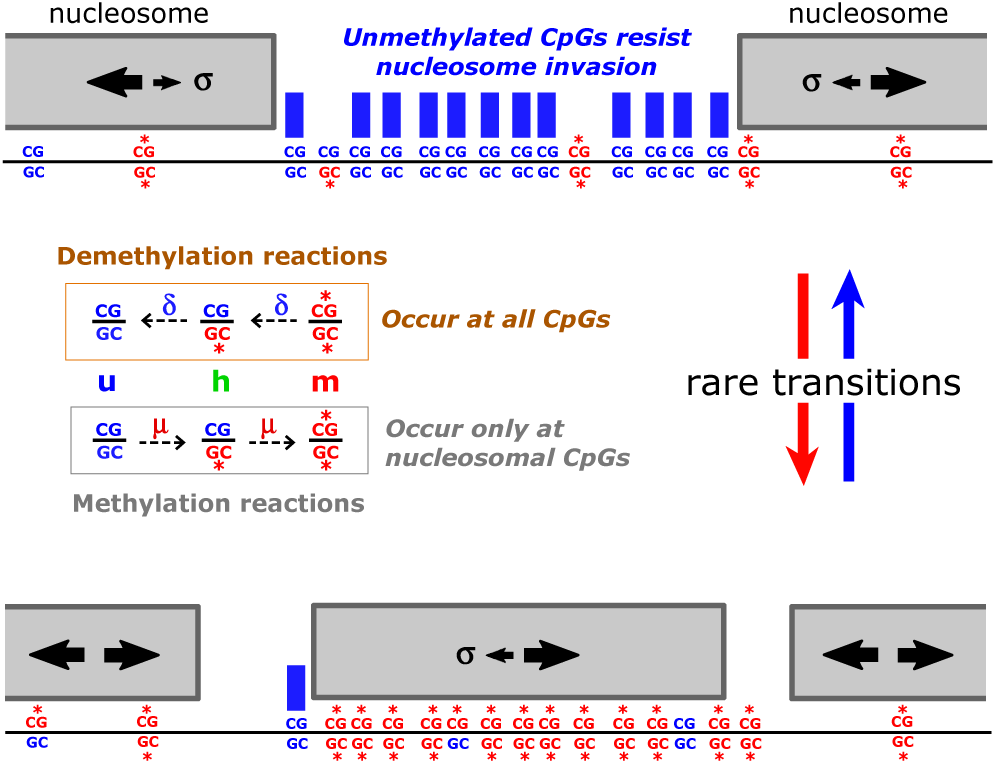
Basic model of nucleosome-DNA methylation interaction. Nucleosomes perform random walks along the DNA, excluding each other and also being restricted by unmethylated CpG sites (u). The restriction in movement only acts from u sites at the boundary of the nucleosome, as it is assumed to reflect binding of some occluding proteins. Demethylation of hemimethylated CpG sites (h - one DNA strand methylated) or fully methylated sites (m - both strands methylated) occurs independently of whether the site is occupied by a nucleosome. In contrast, methylation enzymes are assumed to act only when the CpG site is covered by a nucleosome. The figure shows a dense cluster of CpG sites that can be mostly unmethylated and nucleosome-free, or alternately can be mostly methylated covered by a nucleosome.

In the basic model (Model A) the following actions take place in each time step:

- [Nucleosome movements] This is performed *N(t)* times: One selects a random nucleosome and attempts to move it +2 bp or −2 bp along the DNA. If the attempted nucleosome move brings its center within 144 bp from the center of a neighbor (edges overlapping), the movement is aborted. Unmethylated CpG sites are assumed to be able to bind proteins that exclude nucleosomes, thus if the suggested nucleosome movement brings it on top of an unmethylated CpG site, the move is accepted with probability *σ*, otherwise the move is aborted. Movement over a methylated CpG is always accepted.
- [Nucleosome insertion] For each DNA location where there is > 400 bp between nucleosome centers, or > 250 bp between the boundary and the nearest nucleosome center a nucleosome insertion attempt is made with probability 0.001. A random position for insertion is chosen from those that are > 144 bp from the center of a neighbor. For every unmethylated CpG site within 72 bp of the position, the probability of insertion is reduced by a factor *σ.* The insertion rate is set to correspond to once every time an individual nucleosome has performed about 1000 random steps of size ± 2 bp. Thus insertions occur at a timescale comparable to typical displacement of one nucleosome moving across its own diameter. The results reported in this paper do not depend on this rate, but the parameter values at which the transitions happen will change with nucleosome insertion rate.
- [Methylation] With probability *μ* · *n*_*cpg*_ one selects a random CpG site and attempts to add a methyl group to this site. This move can only be performed if the CpG site is bound to a nucleosome (Fig 1), that is, if the site is within 72 bp from a nucleosome center. If the CpG site is u, then it is changed to h. If the site is h it is changed to m. An m site remains unchanged.
- [Demethylation] All CpGs are subject to demethylation irrespective of whether they are nucleosome bound (Fig 1). With probability *δ · n*_*cpg*_ one selects a random CpG site and attempts to demethylate it. If the site is m it is converted to h. If the site is h it is coverted to u. For a u sites there is no change.

Apart from specifying the system dimensions, the model contains 3 important parameters. One is *σ* that depends on the probability that an unmethylated CpG site will be covered by a DNA binding protein that prevents the nucleosome from making its attempted move. A value of σ = 1 corresponds to nucleosome movement that is independent of the underlying methylation status of the CpG sites, whereas smaller *σ* parameterize increasing occlusion strength. In most simulations shown we use *σ* = 0.1.

The model’s assumptions that nucleosome occupation is resisted by unmethylated CpG sites but not by methylated and hemimethylated CpGs, has some experimental support. Unmethylated CpGs are recognized by ZF-CXXC domain proteins, such as Cfp1 and KDM2A, which bind to unmethylated CpG islands [34–36]. KDM2A has been shown to not bind to nucleosomal DNA, probably because the ZF-CXXC domain and histones make competing DNA contacts [36]. Occupation of DNA by nucleosomes and by sequence-specific binding proteins, such as transcription factors, is generally mutually exclusive for similar reasons [25, 37, 38]. In contrast, the methyl-CpG binding domain (MBD), used by the major class of proteins that recognize methylated CpG sites [39], *is* able to bind nucleosomal DNA, as it contacts only one side of the DNA helix [40].

The two parameters *μ* and *δ* quantify the rate at which methylation and demethylation enzymes act on the CpG site. For simplicity, we set equal rates for the u→h and the h→*m* methylations and equal rates for the m→h and h→u demethylations. The ratio *δ/μ* is the relative strength of the opposing reactions, whereas their absolute size sets the rate of CpG conversions relative to nucleosome movements. As long as *μ* and *δ* are less then 0.001 (i.e much slower than nucleosome movement), then their absolute values do not affect our simulations. The basic model thereby reduces to a two parameter model, and we in this paper assume a fixed value of *μ* = 0.0001 and vary *δ*.

There is currently no direct experimental evidence to support the model’s assumption that methylation of unmethylated or hemimethylated CpGs requires them to be nucleosome-bound, though this would be a simple explanation of the observed correlation between nucleosome occupation and DNA methylation [26–28]. In vitro experiments with DNA methyltransferases DNMT1, DNMT3A and DNMT3B are somewhat inconsistent, but do not clearly indicate any improved methylation for nucleosomal DNA over naked DNA [41, 42]. However, additional factors are likely to affect DNA methylase targeting in vivo. For example, DNMT3L interacts with The H3 histone tail and recruits DNMT3A [43], and DNMT3A and DNMT3B are anchored to DNA-methylated nucleosomes in vivo [44], though it is not clear whether these nucleosome-bound enzymes act within the nucleosomes or on nearby non-nucleosomal DNA.

We also explored two processes that give additional cooperativity between either u sites or m sites.

- [Nucleosome movement: u-u cooperativity (Models B, C and D)] Here we introduce cooperativity in nucleosome exclusion by assuming that pairs of u sites can be bound cooperatively by nucleosome-excluding proteins. When a nucleosome attempts to move onto a u site, the move is prevented if the attacked u site is in protein-mediated contact with another u site. This is determined by randomly sampling *v* random DNA sites at distance *d* from the attacked u site, each with probability *1/d* (i.e. representing the decay in contact probability over distance with a power law with exponent −1). If at least one of these *v* DNA sites is a u, then the attacked u is considered cooperatively bound and the nucleosome movement is blocked. Note that this distance-dependent u-u cooperativity is different from the nearest-neighbor cooperativity used in [45].
- [Methylation recruitment: m-m collaboration (Models C and D)] Nucleosome-bound methylated CpG sites are assumed to recruit enzymes that can methylate other nucleosome-bound CpG sites. This step is performed *n*_*cpg*_ times: With probability *ω* one selects a random CpG site and attempts to let it recruit an enzyme to methylate another (target) CpG site. If the selected site is not fully methylated, or is not covered by a nucleosome then the recruitment attempt is aborted. The target site is chosen by selecting a random distance *d* from the recruiting site with probability *1/d.* If the target site is not a CpG, or if it is not covered by a nucleosome, the attempt is aborted. If the target site is u, then its state is changed to h. If the target site is h, it is changed to m. If the target site is m, then there is no change. The recruitment is different from our earlier models [8, 22, 45] in that it is nucleosome based and only incorporates recruitment acting from fully methylated CpG sites.

In the implementation of u-u cooperativity a higher value of *v* corresponds to a higher probability of forming a DNA loop with one of the u-sites along the chromosome. The *v* attempts in the u-u cooperativity step can completely replace the parameter *σ* in the simple model (Models B and C), but can also be used to complement a single site repulsion factor *σ* < 1 (Model D). Each of the *v* attempts decreases the chance for a nucleosome to invade a given *u* site, and a high *v* accordingly corresponds to a low value of *σ.* A sampling number *v* ~ 25 provides about the same resistance to invasion of an *N* = 15 cluster as *σ* = 0.1 when only sampling loop distances that are less than 3200 base pairs of DNA.

The rate of methylation recruitment attempts (Models C and D) is chosen to be 100 times the rate of passive methylation attempts (*ω* = 100 · *μ*). Note that *ω* is an apparent rate because many of the selected sites for potential methylation are not CpG sites. In practice there is a very high abortion rate for this step, which is also witnessed by the fact that adding a recruitment with *ω* = 100 · *μ* is counterbalanced by only a factor 2.5 fold increase in de-methylation reactions (*δ* = 0.4 → 1) when comparing Models B and C.

Although we do not incorporate DNA replication in the model, we believe that this does not significantly lessen the relevance of the model to real cells. DNA replication has the effect of giving occasional mass conversions of all m’s to h’s and half the h’s to u’s, which without compensating mechanisms would destabilize the hyper-methylated state. However, maintenance methylation enyzymes, catalysing the h to m reaction, are thought to be highly active at the DNA replication fork [7] and should minimize this effect. Conversion of h to u by DNA replication could be compensated for in the model by decreasing the standard m to u and h to u conversion rates. DNA replication also results in histone displacement and incorporation of new histones, altering the locations of nucleosomes [46]. However, there appear to be cellular mechanisms that quickly re-establish dense nucleosome packing [32], and as long as the pre-existing and newly inserted nucleosomes avoid high-u regions, this effect of DNA replication should also be minimal. Techniques such as fluorescence recovery after photobleaching (FRAP) generally indicate fast binding rates for transcription factors [47, 48], so we expect inhibition of nucleosome binding to be rapidly re-established after replication.

## III. RESULTS

Fig. 2 illustrates the basic model (Model A) applied to CpG clusters of various sizes and densities. The parameters *σ* and *δ/μ* are chosen such that a cluster of 15 CpG sites is about equally likely be methylated as to be unmethylated. The CpG sites in the cluster are set to be 10 bp from each other, while CpG sites outside the cluster are 100 bp from each other. These distances are typical of those seen in the human genome [22]. Noticeably, the model produces alternating periods where the cluster is unmethylated (blue), or highly methylated (red). The model also predicts that periods of low methylation are associated with exclusion of nucleosomes (grey dots) from the cluster. Around unmethylated clusters, the model reproduces the phasing of nucleosomes seen next to nucleosome-excluding regions on cellular DNA [31].

**FIG. 2.**
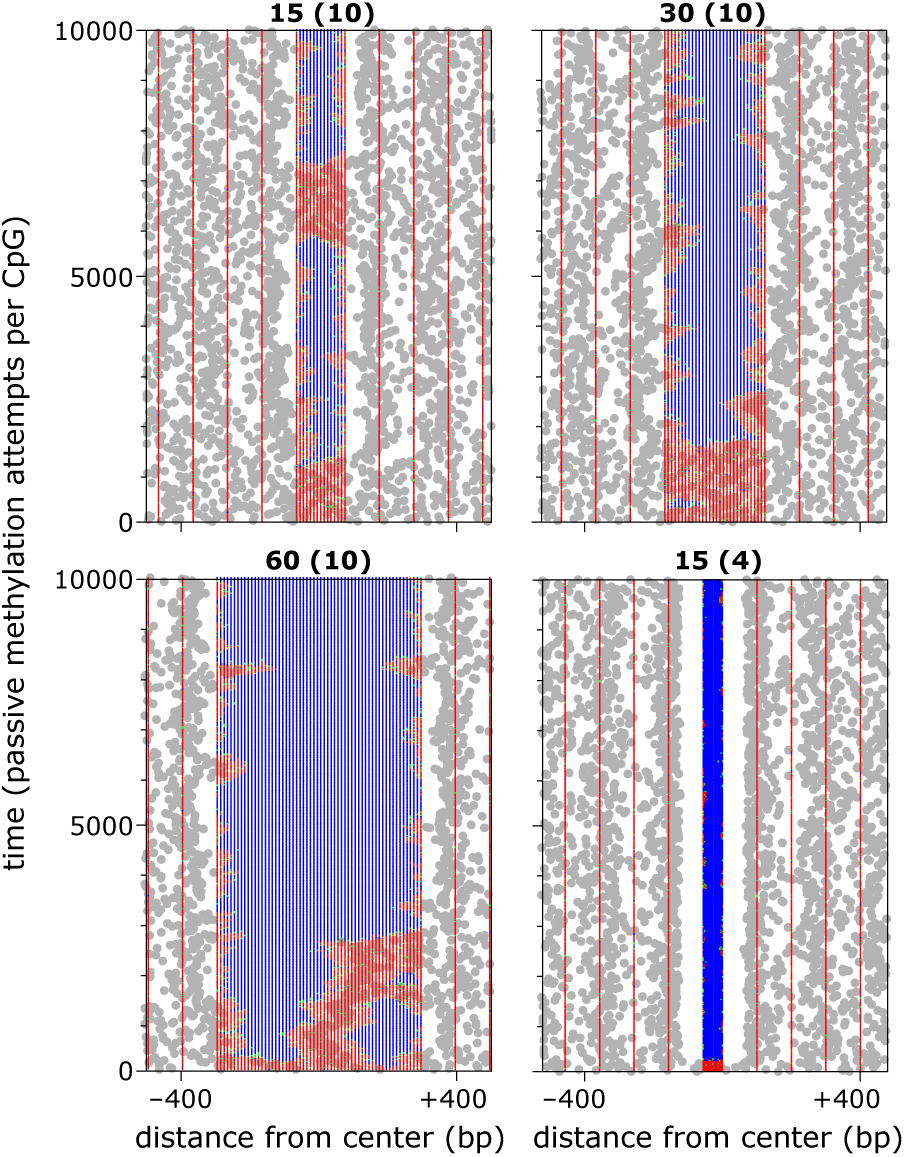
Simulation of the basic model (Model A) *μ* = 0.0001 and *δ/μ* = 0.06 using a L=3200 bp of DNA with a central CpG cluster and 100 bp distances between CpG sites outside the cluster. Grey dots mark the center position of each nucleosome (each nucleosome covers 72 bp on each side). Red, green and blue dots mark m, h and sites, respectively. Numbers on top of each panel refer to the cluster size in number of CpGs, with the spacing in bp (center to center) between the CpG sites in the cluster in parentheses.

As reported recently, [22] the behaviour of CpG clusters in the human genome depends systematically on their size and density. Sampling across cell types, it was found that clusters with less than 10 CpGs tend to be hyper-methylated, clusters with 15-30 CpGs have methylation status that are bimodally distributed, whereas large clusters systematically tend to be unmethylated. Also it was found that this overall behaviour is modulated by the average distance between CpG sites in the cluster, with denser clusters being more likely to be un-methylated. These overall trends are all reproduced by our simple model, as seen in Fig 2 and the upper section of Fig 3.

**FIG. 3.**
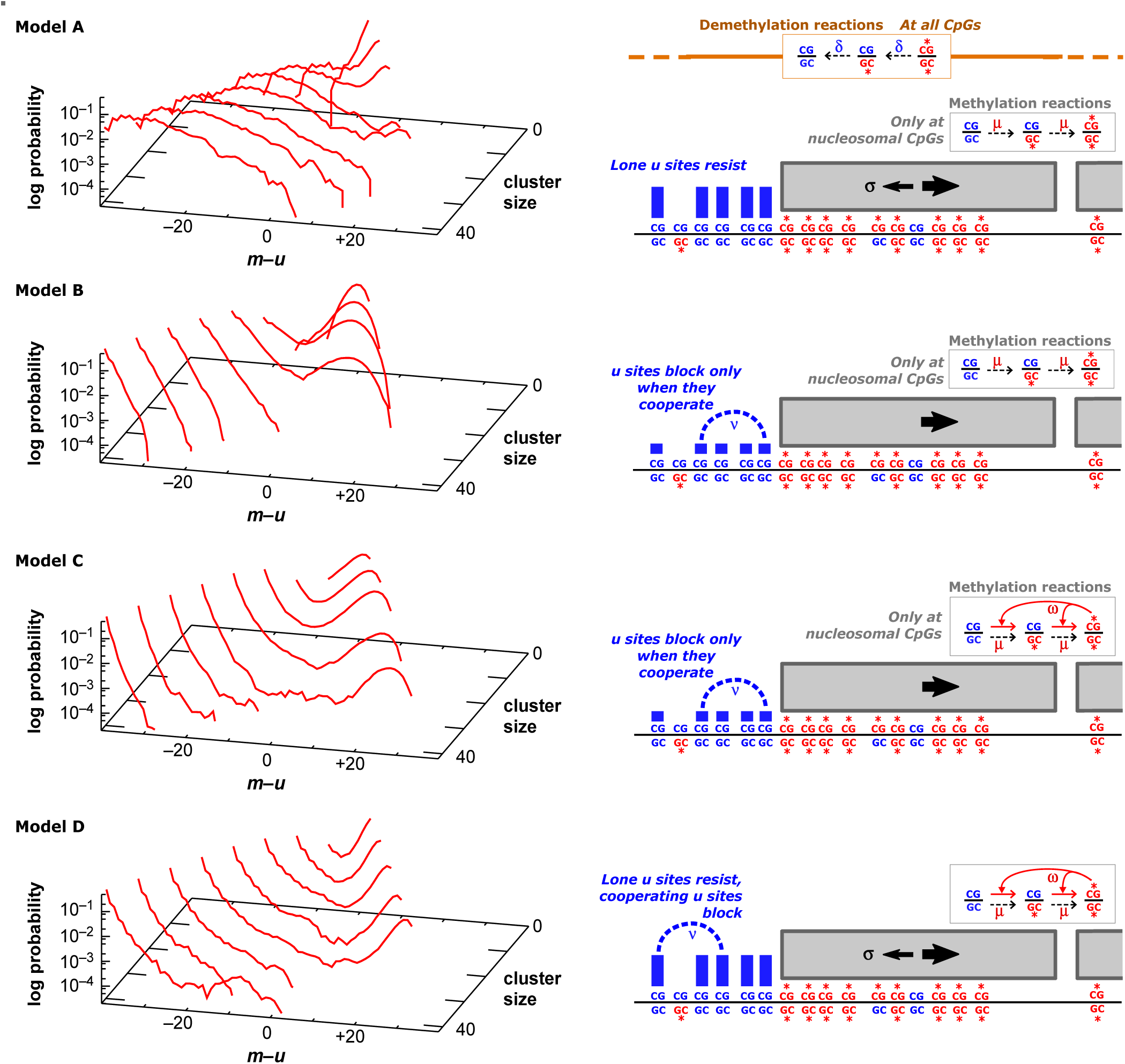
Time averaged distributions of the number of methylated CpG sites minus number of unmethylated sites in the cluster for the four models discussed in this paper. Simulations all use *μ* = 0.0001 and consider clusters positioned in the center of a 3200 bp segment of DNA, with 10 bp between CpG sites within the cluster and 100 bp between CpGs outside the cluster. Model A is the basic model with *δ/μ*, = 0.06 and *σ* = 0.1. Model B is model A with u-u cooperativity added, with *δ/μ*, = 0.4, *v* = 25 and *σ* = 1 (a single u site does not inhibit nucleosomes). Model C is as Model B but also includes methylation recruitment cooperativity, with *δ/μ*, = 1 and *ω/μ*, = 100. Model D is a hybrid between C and A, using *δ/μ*., = 0.4, *ω/μ*, = 100, *v* = 15 and *σ* = 0.3 (single u site inhibits nucleosomes).

This simple model provides positive feedback through a double-negative loop: (1) a u site resists nucleosome occupation, and (2) nucleosome occupation allows a u site to be lost through methylation. This feedback is able to produce bimodality because a nucleosome is large enough to cover multiple CpG sites, giving implicit cooperativity between unmethylated sites in the same way as envisioned in Widom’s classical papers on implicit cooperativity between transcription factor binding to sites that can be occluded by the same nucleosome [49, 50]. When a nucleosome is positioned within parts of the cluster, the covered sites are constantly being methylated, which in turn allows the nucleosome to move back and forth across the cluster, allowing remethylation of CpG sites that are occasionally demethylated. On the other hand, when the CpG cluster is unmethylated, the nucleosome rarely enters the cluster because it has to move against the bias created by multiple unmethylated sites.

We examined the degree of cooperativity of our model as parameterized by the minimal number of unmethylated CpG sites that can facilitate a transition, and found that the presence of 2-3 unmethylated CpG sites within a cluster makes a transition to the unmethylated state likely. Therefore, in terms of actual demethylation reactions the bistability of the system is weak, as shown by the absence of a deep valley in the Model A *m*-*u* histogram in Fig. 3. We therefore considered additional mechanisms for maintaining stable epigenetic states of an intermediate sized CpG cluster.

One possibility is to further explore the idea of nucleosome exclusion by binding of proteins to unmethylated CpG regions. In the basic model this is implemented by a simple factor *σ*, representing the likelihood that a single u site is bound by a nucleosome-excluding protein. However, such proteins may also cooperate by direct protein-protein interactions and thereby exclude nucleosomes more strongly when there are multiple sites. The unmethylated-CpG binding protein KDM2A does not bind low CpG density DNA even when these sites are unmethylated [36], implying that multiple nearby u sites cooperate to provide strong binding. In addition, the ZF-CXXC domain protein Cfp1 recruits enzymes that generate H3K4 methylation [34], a histone modification associated with active promoters and transcription factor recruitment. Indeed, many unmethylated DNA regions bound by ZF-CXXC proteins show promoter activity and bind RNA polymerase [35]. Thus, larger unmethylated clusters are likely to bind a variety of interacting proteins.

We here assume that such protein cooperativity may occur at any distance, but with less probability with increased separation on the DNA (see methods). From the histograms for Model B in Fig 3, we see that such a model easily provides large bistability for size 15 clusters, while preserving the hyper-methylation of small clusters and low methylation of large clusters seen with the simple model.

However, Models A and B do not reproduce two other noteworthy features of CpG clusters in human methylomes [22]. First, hyper-methylated clusters show a higher level of methylation (~0.9) than the general ‘background’ low density CpG sites (~0.8). Second, the methylation level of low density CpG sites near hyper-methylated clusters is elevated above these background methylation levels. This increased methylation decreases with distance from the cluster but extends for considerable distances, with methylation levels still halfway between the cluster level and background levels as far as 30 CpG sites ('~2500 bp) from the cluster. These features suggest some long-range methylation-promoting effects of methylated CpGs [22].

In our previous models of CpG methylation we proposed long-range collaboration between CpG sites, where methylated CpGs recruit enzymes that methylate CpGs in the surrounding DNA [8, 22]. This positive feedback, capable of acting over longer distances, should increase cluster methylation and cause the cluster to increase methylation of surrounding low density CpG sites. It should also aid bistability. Such collaboration has some experimental support. The methyl-CpG binding protein MeCP2 can recruit DNMT1 [51], potentially allowing m sites to stimulate methylation of nearby sites. Collaboration may be more indirect, and utilize histone modifications. For example, MeCP2 can recruit enzymes that methylate histone H3K9 [52], which in turn can recruit proteins that interact with DNMT1 and DNMT3A [53]. Following [22] we represent these reactions in terms of a simple direct recruitment process, as described by the m-m cooperativity in the methods section.

Model C is the same as Model B but with the addi-ton of long range collaborative methylation of moderate strength. We also tested a variant of Model C where nucleosome exclusion is not completely dependent on u-u cooperation *(σ* < 1; Fig 3, Model D). These models preserve the cluster size and density dependence of Model B, and provide bistability for intermediate sized clusters, as well as extending the range of cluster sizes that are bistable (Fig 3). Fig 4 (upper panels) shows that Model D can generate robust bistability and strong nucleosome phasing. In these models, hyper-methylated clusters also show increased methylation relative to the low-density CpG background methylation. Model D gave methylation levels for hyper-methylated clusters and low-density CpG regions of 〈m + *h/2〉* ~ 0.9 and ~ 0.8, respectively, matching observed values. Methylation levels also decreased steadily with increasing distance from the cluster (Fig 4, lower panel). However, we note that Models C and D do not fully account for the observation that increased methylation can extend for several thousands of base pairs around certain methylated CpG clusters [22].

**FIG. 4.**
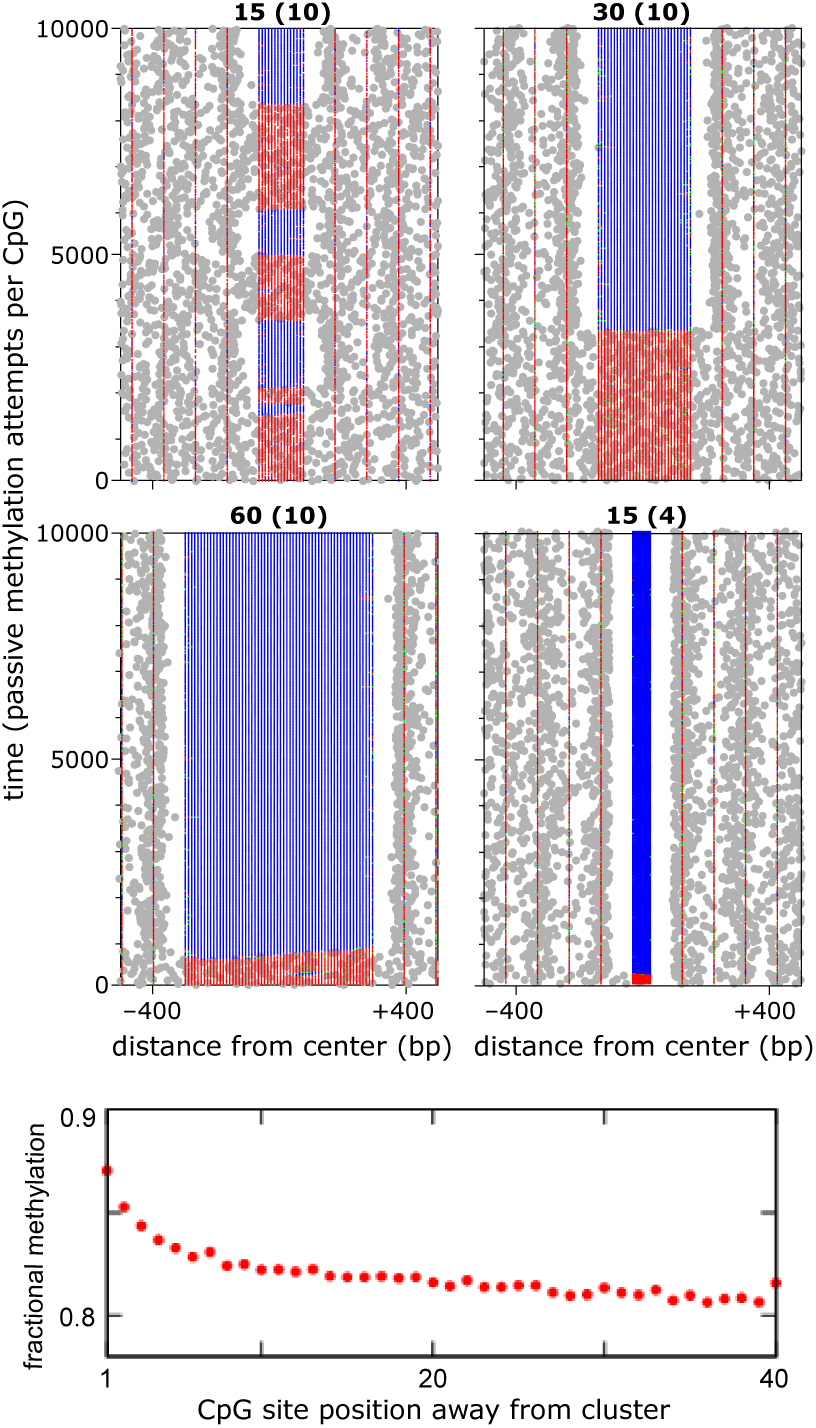
Simulations of Model D (see Fig 3) for clusters of different sizes and densities. Simulations all use *μ* = 0.0001, *δ/μ* = 0.4, *ω* = 100, *v* = 15 and *σ* = 0.3 (single u site inhibits nucleosomes), and consider clusters positioned in the center of a 3200 bp segment of DNA, with 10 bp between CpG sites within the cluster and 100 bp between CpGs outside the cluster. The upper four plots are as in Fig 2. The bottom panel shows the methylation levels (〈*m* + *h/2〉)* for CpGs around a hypermethylated cluster of 15 CpG sites with 10 bp separation in Model D. Methylation near the cluster is elevated compared to the methylation far from the cluster.

## IV. DISCUSSION

We propose and investigate detectable consequences of a feedback between nucleosome positioning and the methylation status of local CpG sites. On a large scale the system consists of nucleosomes that perform a random walk and at the same time modify the substrate that they walk on. The central idea is that one “walker” initially has difficulty in entering a “resistant” region, but once it does so it changes the properties of the region to facilitate entry of more walkers. This in itself is a positive feedback, but it can only provide a gradual transition as one changes the parameters that govern the entry of walkers into the region. However, nucleosomes are in addition extended objects that cover many CpG sites at the same time. It is this “Widom” cooperativity [49, 50] that provides the relatively long lifetimes of the alternative states of the medium size cluster shown in Fig. 2.

The simple model reproduces patterns found for CpG clusters in mammals, including bimodality for intermediate sized clusters, persistent hyper-methylation of small clusters, and persistent low methylation of large clusters (Fig. 2 and Fig. 3A). Also it correctly predicts that dense clusters are more unmethylated (last panel in Fig. 2). These features were also reproduced by the model of [22], but required the use of two somewhat arbitrary and different functions to represent the decay of interaction probability between CpGs as the DNA distance between them increases. In that model, the effect of recruited methylation enzymes was assumed to decay slowly over distance as a power law but with an offset of ~200 bp, while the effect of recruited demethylation enzymes was assumed to be more short range and decay exponentially with a scale of ~170 bp [22]. In the current model, these intermediate distance scales are simply set by the size of the nucleosomes, and we entirely avoided a requirement for recruited de-methylation reactions.

However, the simple model alone predicts quite long lasting transitions between alternative cluster states and thus does not provide a clear separation between the extremes of methylation. We therefore explored additional mechanisms for cooperativity between CpG sites.

The extended models all included resistance to nucleosome occupation by explicit long range cooperativity between u sites. This addition gave increased bistability, even without positive feedback through recruitment of DNA methylation (Model B), while retaining the dependence of methylation on cluster size.

Addition of DNA methylation recruitment (Models C and D) increased the range of cluster sizes that were capable of bistability and gave a better match to observed methylation levels within and around methylated clusters. Also, both models could reproduce an elevated methylation seen for low density CpGs around a methylated cluster and its decrease with distance from the cluster. Neither of these models was however able to account for the full spatial extension of the elevated methylation levels around hypermethylated CpG islands, pos-sibily hinting at a recruited methylation reaction that is of longer range than the 1/*d* assumed in this paper.

The key mechanisms in our model are exclusion of nucleosome occupation by unmethylated DNA, and strong coincidence between DNA methylation activity and CpG positioning on a nucleosome. We note that experimental evidence for the second mechanism is currently lacking, and the actual forces that are at play around CpG clusters are still largely unknown. Real CpG clusters are exposed to various factors that are not determined solely by their size and density, including dynamic sequence-specific binding of transcription factors and associated histone modifiers and nucleosome remodelers. However, the fact that genome-wide statistics of the state of CpG clusters show systematic features as a function of simple static characteristics strongly suggests that methylation is not just a downstream signal from transcription factors that govern gene activity. The models proposed here address these overall features, and utilizing known or plausible mechanisms, can reproduce most of the observed general features of DNA methylation in terms of rules that are applied uniformly to all CpG clusters.

## ACKNOWLEDGMENTS

IBD acknowledges support from the Australian National Health and Medical Research Council (NHMRC grants GNT1025549 and GNT1025549). We thank Jan Haerter for careful reading of the manuscript.

## References

[1] Jaenisch, R. and Bird, A. (2003) Epigenetic regulation of gene expression: how the genome integrates intrinsic and environmental signals. Nature genetics, 33, 245–254.

[2] Bird, A. (2007) Perceptions of epigenetics. Nature, 447(7143), 396–398.

[3] Jones, P.A. (2012) Functions of DNA methylation: islands, start sites, gene bodies and beyond. Nature reviews. Genetics, 13, 484–492.

[4] Felsenfeld, G. (2014) A brief history of epigenetics. Cold Spring Harbor perspectives in biology, 6(1), a018200.

[5] Holliday, R. and Pugh, J. E. (1975) DNA modification mechanisms and gene activity during development. Science, 187(4173), 226–232.

[6] Riggs, A. D. (1975) X inactivation, differentiation, and DNA methylation. Cytogenetic and Genome Research, 14(1), 9–25.

[7] Jones, P. A. and Liang, G. (November, 2009) Rethinking how DNA methylation patterns are maintained. Nature reviews. Genetics, 10(11), 805–11.

[8] Haerter, J. O., Lövkvist, C., Dodd, I. B., and Sneppen, K. (2013) Collaboration between CpG sites is needed for stable somatic inheritance of DNA methylation states. Nucleic acids research, p. gkt1235.

[9] Jeltsch, A. and Jurkowska, R. Z. (2014) New concepts in DNA methylation. Trends in biochemical sciences, 39(7), 310–318.

[10] Lorincz, M. C., Schübeler, D., Hutchinson, S. R., Dickerson, D. R., and Groudine, M. (2002) DNA methylation density influences the stability of an epigenetic imprint and Dnmt3a/b-independent de novo methylation. Molecular and cellular biology, 22(21), 7572–7580.

[11] Williams, K., Christensen, J., Pedersen, M. T., Johansen, J. V., Cloos, P. A., Rappsilber, J., and Helin, K. (2011) TET1 and hydroxymethylcytosine in transcription and DNA methylation fidelity. Nature, 473(7347), 343–348.

[12] He, Y.-F., Li, B.-Z., Li, Z., Liu, P., Wang, Y., Tang, Q., Ding, J., Jia, Y., Chen, Z., Li, L., et al. (2011) Tet-mediated formation of 5-carboxylcytosine and its excision by TDG in mammalian DNA. Science, 333(6047), 1303–1307.

[13] Wu, H. and Zhang, Y. (2014) Reversing DNA methylation: mechanisms, genomics, and biological functions. Cell, 156(1), 45–68.

[14] Bhutani, N., Burns, D.M. and Blau, H.M. (2011) DNA demethylation dynamics. Cell, 146, 866–872.

[15] Laird, C. D., Pleasant, N. D., Clark, A. D., Sneeden, J. L., Hassan, K. M. A., Manley, N. C., Vary, J. C., Morgan, T., Hansen, R. S., and Stöger, R. (2004) Hairpin-bisulfite PCR: assessing epigenetic methylation patterns on complementary strands of individual DNA molecules. Proceedings of the National Academy of Sciences USA 101(1), 204–9.

[16] Sontag, L.B., Lorincz, M.C. and Georg Luebeck, E. (2006) Dynamics, stability and inheritance of somatic DNA methylation imprints. J. Theor. Biol., 242, 890–899.

[17] Pfeifer, G.P., Steigerwald, S.D., Hansen, R.S., Gartler, S.M. and Riggs, A.D. (1990) Polymerase chain reaction-aided genomic sequencing of an X chromosome-linked CpG island: methylation patterns suggest clonal inheritance, CpG site autonomy, and an explanation of activity state stability. Proc Natl Acad Sci USA, 87, 8252–8256.

[18] Eckhardt, F., Lewin, J., Cortese, R., Rakyan, V. K., Attwood, J., Burger, M., Burton, J., Cox, T. V., Davies, R., Down, T. A., et al. (2006) DNA methylation profiling of human chromosomes 6, 20 and 22. Nature genetics, 38(12), 1378–1385.

[19] Weber, M., Hellmann, I., Stadler, M. B., Ramos, L., Pääbo, S., Rebhan, M., and Schübeler, D. (2007) Distribution, silencing potential and evolutionary impact of promoter DNA methylation in the human genome. Nature genetics, 39(4), 457–466.

[20] Meissner, A., Mikkelsen, T. S., Gu, H., Wernig, M., Hanna, J., Sivachenko, A., Zhang, X., Bernstein, B. E., Nusbaum, C., Jaffe, D. B., et al. (2008) Genome-scale DNA methylation maps of pluripotent and differentiated cells. Nature, 454(7205), 766–770.

[21] Zhang, Y., Rohde, C., Tierling, S., Jurkowski, T. P., Bock, C., Santacruz, D., Ragozin, S., Reinhardt, R., Groth, M., Walter, J., et al. (2009) DNA methylation analysis of chromosome 21 gene promoters at single base pair and single allele resolution. PLoS genetics, 5(3), e1000438.

[22] Lövkvist, C., Dodd, I.B., Sneppen, K. and Haerter, J.O. (2016) DNA methylation in human epigenome depends on local topology of cpG sites. Nucleic Acids Res, gkw124 [Epub ahead of print].

[23] Cedar, H. and Bergman, Y. (2009) Linking DNA methylation and histone modification: patterns and paradigms. Nature Rev Genet, 10, 295–304.

[24] Felsenfeld, G. and Groudine, M. (2003) Controlling the double helix. Nature, 421, 448–453.

[25] Segal, E. and Widom, J. (2009) What controls nucleosome positions? Trends Genet, 25, 335–343.

[26] Chodavarapu, R.K., Feng, S., Bernatavichute, Y.V., Chen, P.Y., Stroud, H., Yu, Y., Hetzel, J.A., Kuo, F., Kim, J., Cokus, S.J. et al. (2010) Relationship between nucleosome positioning and DNA methylation. Nature, 466, 388–392.

[27] Kelly, T.K., Liu, Y., Lay, F.D., Liang, G., Berman, B.P. and Jones, P.A. (2012) Genome-wide mapping of nucleosome positioning and DNA methylation within individual DNA molecules. Genome Res, 22, 2497–2506.

[28] Portela, A., Liz, J., Nogales, V., Setien, F., Villanueva, A. and Esteller, M. (2013) DNA methylation determines nucleosome occupancy in the 5’-CpG islands of tumor suppressor genes. Oncogene, 32, 5421–5428.

[29] Kornberg R, Stryer L (1988) Statistical distributions of nucleosomes: nonrandom locations by a stochastic mechanism. Nucleic Acids Res, 16, 6677–90.

[30] Möbius, W., and Gerland, U. Quantitative Test of the Barrier Nucleosome Model for Statistical Positioning of Nucleosomes Up-and Downstream of Transcription Start Sites (2010). PLoS Comput Biol, 6, e1000891. http://doi.org/10.1371/journal.pcbi.1000891

[31] Möbius, W., Osberg, B., Tsankov, A.M., Rando, O.J. and Gerland, U. (2013) Toward a unified physical model of nucleosome patterns flanking transcription start sites. Proc Natl Acad Sci USA, 110, 5719–5724.

[32] Osberg, B., Nuebler, J., Korber, P. and Gerland, U. (2014) Replication-guided nucleosome packing and nucleosome breathing expedite the formation of dense arrays. Nucleic Acids Res, 42, 13633–13645.

[33] Clapier, C. R., and Cairns, B. R. (2009). The biology of chromatin remodeling complexes. Annual Review of Biochemistry, 78, 273–304.

[34] Thomson, J. P., Skene, P. J., Selfridge, J., Clouaire, T., Guy, J., Webb, S., Kerr, A. R., Deaton, A., Andrews, R., James, K. D., et al. (2010) CpG islands influence chromatin structure via the CpG-binding protein Cfp1. Nature, 464(7291), 1082–1086.

[35] Illingworth, R.S., Gruenewald-Schneider, U., Webb, S., Kerr, A.R., James, K.D., Turner, D.J., Smith, C., Harrison, D.J., Andrews, R. and Bird, A.P. (2010) Orphan CpG islands identify numerous conserved promoters in the mammalian genome. PLoS Genet, 6, e1001134.

[36] Zhou, J.C., Blackledge, N.P., Farcas, A.M. and Klose, R.J. (2012) Recognition of CpG island chromatin by KDM2A requires direct and specific interaction with linker DNA. Mol Cell Biol, 32, 479–489.

[37] Bai, L., Ondracka, A. and Cross, F.R. (2011) Multiple sequence-specific factors generate the nucleosome-depleted region on CLN2 promoter. Mol Cell, 42, 465–476.

[38] Vierstra, J., Wang, H., John, S., Sandstrom, R. and Stamatoyannopoulos, J.A. (2014) Coupling transcription factor occupancy to nucleosome architecture with DNase-FLASH. Nat Methods, 11, 66–72.

[39] Du, Q., Luu, P.L., Stirzaker, C. and Clark, S.J. (2015) Methyl-CpG-binding domain proteins: readers of the epigenome. Epigenomics, 7, 1051–1073.

[40] Ohki, I., Shimotake, N., Fujita, N., Jee, J., Ikegami, T., Nakao, M. and Shirakawa, M. (2001) Solution structure of the methyl-CpG binding domain of human MBD1 in complex with methylated DNA. Cell, 105, 487–497.

[41] Gowher, H., Stockdale, C.J., Goyal, R., Ferreira, H., Owen-Hughes, T. and Jeltsch, A. (2005) De novo methylation of nucleosomal DNA by the mammalian Dnmt1 and Dnmt3A DNA methyltransferases. Biochemistry, 44, 9899–9904.

[42] Takeshima, H., Suetake, I., Shimahara, H., Ura, K., Tate, S. and Tajima, S. (2006) Distinct DNA methylation activity of Dnmt3a and Dnmt3b towards naked and nucleosomal DNA. J Biochem, 139, 503–515.

[43] Ooi, S.K., Qiu, C., Bernstein, E., Li, K., Jia, D., Yang, Z., Erdjument-Bromage, H., Tempst, P., Lin, S.P., Allis, C.D. et al. (2007) DNMT3L connects unmethylated lysine 4 of histone H3 to de novo methylation of DNA. Nature, 448, 714–717.

[44] Jeong, S., Liang, G., Sharma, S., Lin, J.C., Choi, S.H., Han, H., Yoo, C.B., Egger, G., Yang, A.S. and Jones, P.A. (2009) Selective anchoring of DNA methyltransferases 3A and 3B to nucleosomes containing methylated DNA. Mol Cell Biol, 29, 5366–5376.

[45] Sormani, G., Haerter, J.O., Lövkvist, C. and Sneppen, K. (2016) Stabilization of epigenetic states of CpG islands by local cooperation. Mol BioSyst, DOI: 10.1039/C6MB00044D

[46] Radman-Livaja, M., Verzijlbergen, K.F., Weiner, A., van Welsem, T., Friedman, N., Rando, O.J. and van Leeuwen, F. (2011) Patterns and mechanisms of ancestral histone protein inheritance in budding yeast. PLoS Biol, 9, e1001075.

[47] Phair, R.D., Scaffidi, P., Elbi, C., Vecerova, J., Dey, A., Ozato, K., Brown, D.T., Hager, G., Bustin, M. and Misteli, T. (2004) Global nature of dynamic protein-chromatin interactions in vivo: three-dimensional genome scanning and dynamic interaction networks of chromatin proteins. Mol Cell Biol, 24, 6393–6402.

[48] Mueller, F., Stasevich, T.J., Mazza, D. and McNally, J.G. (2013) Quantifying transcription factor kinetics: at work or at play? Crit Reviews Biochem Molec Biol, 48, 492–514.

[49] Polach, K.J. and Widom, J. (1996) A model for the cooperative binding of eukaryotic regulatory proteins to nucleosomal target sites. J Mol Biol, 258, 800–812.

[50] Miller, J.A. and Widom, J. (2003) Collaborative competition mechanism for gene activation in vivo. Mol Cell Biol, 23, 1623–1632.

[51] Kimura, H. and Shiota, K. (2003) Methyl-CpG-binding protein, MeCP2, is a target molecule for maintenance DNA methyltransferase DNMT1 (2003) J Biol Chem, 278, 4806–4812.

[52] Fuks, F., Hurd, P.J., Wolf, D., Nan, X., Bird, A.P. and Kouzarides, T. (2003) The methyl-CpG-binding protein MeCP2 links DNA methylation to histone methylation. J Biol Chem, 278, 4035–4040.

[53] Fuks, F., Hurd, P.J., Deplus, R. and Kouzarides, T. (2003) The DNA methyltransferases associate with HP1 and the SUV39H1 histone methyltransferase. Nucleic Acids Res, 31, 2305–2312.

